# Two-Photon NAD(P)H-FLIM reveals unperturbed energy metabolism of *Ascaris suum* larvae, in contrast to host macrophages upon artemisinin derivatives exposure

**DOI:** 10.1101/2024.08.30.610452

**Authors:** Zaneta D. Musimbi, Arkadi Kundik, Jürgen Krücken, Anja E. Hauser, Sebastian Rausch, Peter H. Seeberger, Raluca Niesner, Ruth Leben, Susanne Hartmann

## Abstract

Soil-transmitted helminths (STH) are widespread, with *Ascaris lumbricoides* infecting millions globally. Malaria and STH co-infections are common in co-endemic regions. Artemisinin derivatives (ARTs)—artesunate, artemether, and dihydroartemisinin—are standard malaria treatments and are also known to influence the energy metabolism of parasites, tumors, and immune cells. Herein, we explore the potential of ARTs to influence ascariasis either by directly affecting larvae or indirectly by modifying macrophage responses. *Ascaris suum* third-stage larvae and porcine IL-4 polarised (M2-like) macrophages were exposed to ARTs *in vitro*, and their metabolism was evaluated using two-photon NAD(P)H-FLIM. Both larvae and M2-like macrophages exhibited a steady-state bioenergetic profile of high oxidative phosphorylation and low anaerobic glycolysis. In *A. suum* larvae, two metabolically distinct regions were identified, with particularly high DUOX activity in the pharynx compared to the midgut. The metabolic profile of both larval regions were, however, unperturbed by ARTs exposure. In contrast, exposure of M2-like macrophages to ARTs induced a metabolic shift towards high anaerobic glycolysis and reduced metabolic activity, suggesting a possible indirect effect of ARTs on the helminth infection. Overall, two-photon NAD(P)H-FLIM proved to be a powerful tool for studying specific metabolic pathways in *Ascaris* larvae and host macrophages, offering valuable insights into the metabolic mechanisms of drug action on both parasite and host.

## Introduction

Approximately 772–892 million people worldwide are infected with the roundworm, *Ascaris lumbricoides* ^1^; a soil transmitted helminth (STH) prevalent in tropical and sub-tropical regions ^2,3^. Ascariasis involves a hepato-tracheal migration of third-stage larvae (L3) from the intestines, through the caecum and liver to the lungs. The L3 penetrate the alveolar, migrate to the trachea, are expectorated and swallowed back to the small intestine where they molt to L4, L5 and finally adults ^4^. The zoonotic ^5^ roundworm of pigs, *Ascaris suum,* is genetically comparable to *A. lumbricoides* ^6^ and demonstrates similar immune pathology in the pig to that of *A. lumbricoides* in humans. In addition, Meurens *et al* reviewed the close resemblance of pigs to human anatomy, physiology, immune composition among others, making it an ideal model for human infectious diseases ^7^. Thus, *A. suum* infection is a relevant host-nematode model to study human ascariasis.

We showed earlier that migratory *Ascaris* larvae cause tissue injury and type 2 inflammatory infiltration in affected local sites of experimental pigs ^8^ and mice ^9^. Murine pulmonary ascariasis activates alveolar and interstitial macrophage activity ^10^ and induces an IL-4/ IL-5 environment that polarizes resting macrophages (M0 macrophages) into alternatively activated macrophages (M2-like macrophages) ^9,11^. An *in vitro* study by Coakley *et al* found that human blood monocyte-derived macrophages activated by *Ascaris*-infected serum adhered to and immobilized *A. suum* larvae ^12^.

Artemisinin and its semi-synthetic derivatives, dihydroartemisinin, artesunate, arteether and artemether (collectively referred to ARTs herein), are endoperoxide sesquiterpene lactones primarily recognized for their potent antimalarial properties ^13,14^. Due to their effectiveness in rapidly reducing *Plasmodium* parasitemia, World Health Organization (WHO) recommends artemisinin-based combined therapy (ACTs) for treating uncomplicated malaria. Many tropical regions are endemic to both soil-transmitted helminths and *Plasmodium* malaria ^15–17^. A recent field study conducted in a malaria- and schistosomiasis-co-endemic area in Gabon found that ACT treatment for malaria unexpectedly led to the control of urogenital schistosomiasis in co-infected patients ^18^. Additionally, experimental studies on roundworms such as *T. spiralis* ^19^, *Toxocara canis* ^20^ and *Haemonchus contortus* ^21^ demonstrated ART-induced morphological changes *in vitro* and reduced worm burden *in vivo*. Beyond its effects on helminths and malaria, ARTs have shown immunomodulatory properties*. Mancuso* et al reported that artesunate deactivated the IL-4-induced JAK2/STAT3 pathway, leading to reduced expression of CD206, CD16, and CD163, as well as increased nitric oxide (NO) production and the accumulation of inflammatory monocytes ^22^.

The multifunctional properties of ARTs ^19–21,23^ could be attributed to the equally diverse mode of action of the drugs, which include, but not limited to; 1) heme adduct formation, 2) protein alkylation, 3) free radical production ^24^ and, most recently, 4) interference with palmitoylation ^25^. Energy metabolic pathways have also been implicated as both direct and indirect targets of ARTs. Mitochondria are suggested to activate ARTs thereby inducing superoxide radicals that interfere with the electron transport chain leading to ATP depletion ^26–30^. In addition, studies described significantly reduced expression levels of glucose transporter 1 gene and genes encoding glycolytic enzymes in *Plasmodium falciparum* ^31,32^, schistosomes ^33^, cancer cells ^34–37^ and Th17 cells ^38^ after exposure to ARTs.

In this study, we sought to investigate the potential of artemisinin derivatives to impact Ascaris infections, either by directly targeting the larvae or by altering macrophage responses. Given the reported anthelmintic activity and metabolic effects of ARTs, we examined the potential impact of artemisinin, artesunate, and artemether on the energy metabolism of the parasitic nematode, *A. suum* L3, and IL-4 stimulated porcine alveolar-derived macrophages *in vitro*. The ubiquitous metabolic factors NADH and NADPH (hereafter referred to as NAD(P)H) are endogenously fluorescent. Label-free NAD(P)H fluorescence lifetime imaging (FLIM) enables the evaluation of metabolic profiles of living cells and tissues at subcellular level ^39–42^ and has been used to study metabolism under both physiologic and pathologic conditions^43–48^. We have previously adapted and applied NAD(P)H-FLIM to study the metabolism of the murine helminth *Heligmosomoides polygyrus* and of the host intestine *in situ* ^49,50^. Others have applied the method to investigate the germline of live *C. elegans* nematodes ^51^ and also the manipulation of host cell metabolism by *Toxoplasma gondii* ^48^.

In this study, we utilized two-photon microscopy combined with NAD(P)H-FLIM to investigate how ARTs affect energy production and oxidative burst (defense metabolism) in *A. suum* L3 and IL-4 stimulated PAM 3D4/31 cells, a porcine macrophage cell line. Our findings reveal that, under steady-state conditions, both *A. suum* L3 and the macrophages exhibit a metabolic profile characterized by high oxidative phosphorylation (OxPhos) and low anaerobic glycolysis. We further observed that *A. suum* L3 has two metabolically distinct larval regions that remain unaffected by ART exposure. Conversely, M2 macrophages exposed to artesunate exhibited a metabolic shift from high OxPhos/low anaerobic glycolysis to high OxPhos/ high anaerobic glycolysis with a reduction in overall enzymatic activity.

## Materials and Methods

### Drugs and chemicals

Artemisinin (purified as per Zhou *et al.*^52^), artemether (CAS RN 71963-77-4) and artesunate (CAS RN88495-63-0) were provided by the Seeberger laboratory at Max Planck Institute of Colloids and Interfaces (Potsdam, Germany). Drug stock solutions of 100 mM were prepared in 100% DMSO and stored at -20°C.

### *Ascaris suum* egg collection, embryonation and hatching

Adult *A. suum* worms were obtained from the slaughterhouse. All experiments were done in biosafety level 2 laboratories with permission to work with *A. suum* infective eggs, larvae and adults (S2-919/94).

*Ascaris suum* eggs were collected from *A. suum* female adults obtained from a slaughterhouse, prepared and cultured^53^. Briefly, the female adults were separated from males and washed with 0.9% physiological NaCl. Thereafter, they were incubated for three days in balanced salt solution (HBSS-AB: 127 mM NaCl, 7.5 mM NaHCO3, 5 mM KCl, 1 mM CaCl2, 1 mM MgCl2), 0.1% glucose,

200 U/ml penicillin, 200 μg/ml streptomycin, 2.5 μg/ml amphotericin and 50 μg/ml gentamycin (PAN-Biotech GmbH, Germany) (referred to as medium hereafter) at an atmosphere of 37 °C, 5% CO_2_. *Ascaris suum* eggs were collected every 24 hours whilst changing the media. The harvested eggs were washed and incubated at 30 °C for embryonation. Fully embryonated eggs were washed and separated from un-fertilized and un-embryonated eggs using a sucrose step gradient of 18, 20 and 23% (w/v)^54^. Embryonated *A. suum* eggs (recovered from 23% sucrose layer) were then hatched using the glass bead method^55^ with slight modifications. Briefly, eggs in medium were added into an Erlenmeyer flask containing glass beads and hatching occured in an incubator shaker at 37 °C, 100 rpm. To separate the hatched *A. suum* larvae from eggshells or unhatched larvae, content in the Erlenmeyer flask was washed, transferred onto a 30 μM sieve in 100 ml beaker containing 10 ml HBSS media and incubated at 37 °C, 5% CO_2_ for a minimum of three hours and a maximum of 15 hours to allow for migration. Migrated larvae were washed and immediately used for drug exposures.

In 96 well plates, *A. suum* (∼1000 L3/well) in media were exposed to artesunate and artemether at concentrations of 20- and 200 μM for 24 hours and 200 µM for 48 hours at 37 °C and 5% CO_2_ with vehicle and negative controls run in parallel. Worms were washed three times with fresh medium and transferred into 1.5 ml microcentrifugation tubes. Worms were transferred to glass slides and immobilized by NemaGel (In Vivo Biosystems, Eugene, Oregon, USA). NemaGel was used to trap the worms for imaging without injuring them.

### Porcine alveolar-derived macrophages

Porcine alveolar-derived macrophages (PAM) 3D4/31 cell line ^56^ (provided by Dr. Karsten Tedin, Institute of Microbiology and Epizootics, Freie Universität Berlin) were cultured as monolayers in complete Iscove’s Modified Dulbecco’s Medium (IMDM (PAN-Biotech GmbH, Germany) supplemented with 10% heat-inactivated fetal calf serum, 1% penicillin/streptomycin; herein referred to as CIMDM) at standard tissue culture conditions (37 C, 5% CO2). Experiments were performed within five passages (passage 26-30) after seeding the original frozen stock. In 6 well plates, the porcine macrophages (2 × 10^6^ cells/well) were pre-stimulated overnight with either 1µg/ml LPS/1µg/ml IFNy or 50ng/ml IL-4. Stimulated and unstimulated cells were exposed to 1µM artesunate and incubated for 24 hours at 37°C and 5% CO_2_ with vehicle and negative controls run in parallel. Experiments were done at ∼90% cell confluency and in duplicate. For imaging, CIMDM was aspirated and replaced with 2 ml Dulbecco’s phosphate-buffered saline DPBS.

### Two-photon microscopy FLIM

The two-photon FLIM experiments were carried out on a specialized commercial laser scanning microscope (TriMScope II, LaVision Biotec, Miltenyi, Bielefeld, Germany), as previously described^57^ with a Titanium: Sapphire laser (80 MHz, 140 fs pulses, Chameleon Ultra II, Coherent, Duisburg, Germany) tuned at 765 nm for NAD(P)H excitation. The beam was focused into the sample by a water-immersion objective (20×, NA 1.05, Apochromat, Olympus, Hamburg Germany). To ensure the selective detection of NAD(P)H fluorescence, the emitted signal was collected in the range 466±20 nm, excluding other endogenous contributions ^58–61^. A GaAsP PMT (Hamamatsu, Herrsching, Germany) connected to a time-correlated single photon counting device (TCSPC LaVision Biotec, Miltenyi, Bielefeld, Germany) was used detect fluorescence (55 ps bin width, over at least 9 ns). The acquired images were typically 200×200 µm² for the macrophages and 245 × 245 µm for the larvae (both at 512 × 512 pixels), the pixel dwell time was 5877 ms. Five frames were acquired per field of view (FoV) in fast mode and subsequently added together in order to increase the pixel dwell time (actual image time 29.4 s) and thus the signal-to-noise ratio.

The photon arrival histograms were subsequently added up before analyzing.

### NAD(P)H-dependent enzymes used for the phasor analysis of NAD(P)H-FLIM data

Transcriptome analysis of mammalian cells was used to obtain thirteen most abundant NAD(P)H-dependent enzymes as detailed in Table 1. These were used for the phasor analyses of the porcine macrophages data (see next paragraph). Before this analytical system was applied to *A. suum*, the expression of these enzymes in *A. suum* was verified using the genome database for parasitic worms “WormBase ParaSite” (https://parasite.wormbase.org/index.html, last view 16.08.2023; original source of the *Ascaris* data set^62^) and the proteins confirmed via mass spectrometry of *A. suum* L3. The protein sequences were obtained from the UniProt database^63^ and blasted against the respective human enzymes. The InterPro database^64^ was thereafter used to determine the homology of the NAD(P)H binding sites. The system was adapted accordingly (Table 1). Since no DUOX enzyme was annotated in the *A. suum* genome, the human DUOX enzyme sequence was blasted against *A. suum* genome. The resulting unannotated sequence was blasted against *Caenorhabditis elegans,* and homology of NAD(P)H determined. It was further referred to as DUOX-like enzyme in the text.

**Table 1.** Abundant NAD(P)H-dependent enzymes present in mammalian cells and *A. suum* L3.

| Abbreviation | Full name | Fluorescence lifetime of NAD(P)H [ps] | Mammalian cell pathways involved | Expression in *A. suum* L3 |
| --- | --- | --- | --- | --- |
| nan | pixel could not be allocated | - | - | - |
| Free NAD(P)H | Unbound NAD(P)H | 450 | Inactive metabolism | ✓ |
| MDH | Malate dehydrogenase | 1240 | TCA cycle | ✓ |
| HADH | Hydroxy acyl-coenzyme-A dehydrogenase | 1360 | Beta oxidation | ✓ |
| LDH | Lactate dehydrogenase | 1600 | Anaerobic glycolysis/fermentation | ✓ |
| G6PDH | Glucose-6-phosphate dehydrogenase | 2005 | Pentose phosphate pathway | x |
| SDH (NADH) | Sorbitol dehydrogenase (bound to NADH) | 2010 | Fructose synthesis | x |
| GAPDH | Glyceraldehyde-3-phosphate dehydrogenase | 2050 | Glycolysis | ✓ |
| IDH | Isocitrate dehydrogenase | 2170 | TCA cycle | ✓ |
| SDH (NADPH) | Sorbitol dehydrogenase (bound to NADPH) | 2260 | Fructose synthesis | x |
| PDH / CTBP1 | Pyruvate dehydrogenase / C-terminal binding protein 1 (nuclear) | 2470 | Acetyl-CoA synthesis/gene silencing | ✓ (PDH)<br>x (CTBP1)) |
| iNOS | Inducible nitric oxide synthase | 2550 | Immune defense | x |
| ADH | Alcohol dehydrogenase | 2600 | Detoxification | ✓ |
| NOX/DUOX | Activated NADPH oxidation | 3650 | Oxidative burst | ✓ DUOX<br>x NOX |
| ✓ “is expressed”; x “is not expressed” |  |  |  |  |

The time-domain fluorescence lifetime data were analyzed by an algorithm written in Python (Version 3.9.7)^57^. In brief, the add-up photon arrival histogram was transformed into the normalized phase domain by computing the discrete Fourier transformation, resulting in a complex number consisting of a real and imaginary part which give the coordinates in a phasor plot^65^ (Supplementary Fig. S1). The half circle in the normalized phasor plot represents all possible mono-exponential fluorescence lifetimes and limits the range that can be used for analysis. By adequate signal-to-noise ratio (SNR > 5)^66^ each pixel in the phasor plot was assigned to one of the ten most abundant NAD(P)H-dependent enzymes found in *A. suum*. Here we achieved an SNR of ∼14 (three time higher than the minimum), which corresponds to a mean photon count per pixel of ∼30 photons/pixel or approximately ∼35000 photons/cell (min. 4277 photons/cell or larvae), ensuring the quality of our FLIM measurement.

After assigning each pixel in the images to one of the NAD(P)H-dependent enzymes or unbound NAD(P)H, these pixels were transferred back to space domain to create an enzyme map (Supplementary Fig. S1). In the phasor plot, the length of the vector pointing from free/unbound NAD(P)H “free” towards the position of the lifetime of the assigned enzyme gives the enzyme-bound NAD(P)H activity in each pixel of the image (Supplementary Fig. S1). Pixels with a fluorescence lifetime of 0.45 ns indicate unbound NAD(P)H and are interpreted to have a metabolically inactive status, since no NAD(P)H-dependent metabolic process is catalyzed. Enzymes dominant in anaerobic glycolysis/ β-oxidation of fatty acids are described to have shorter fluorescence lifetimes ranging between 1-2.26 ns whilst enzymes dominant within aerobic glycolysis/oxidative phosphorylation (OxPhos) are described to have longer fluorescence lifetimes ranging between 2.47-2.6 ns. Therefore, pixels with fluorescence lifetimes of 1.24 – 2.6 ns are collectively classified as being active in energy metabolism. NAD(P)H bound to NADPH oxidases have been described as having a very long fluorescence lifetime of 3.65 ns^67^.

### Segmentation and FLIM image analyses

#### 1. Porcine macrophage cell line

Each NAD(P)H-FLIM image of the porcine macrophages contained approximately 125 cells. To display the heterogeneity of the single cells, we segmented the cells with Cellpose ^68^ obtaining approximately 1250 cells (from 10 fields of view) per treatment. We applied the resulting masks to the corresponding enzyme and activity maps^66^. A metabolic state of inactivity was denoted by free NAD(P)H. An active energy production metabolic state was considered when NAD(P)H was bound to the enzymes MDH, HADH, LDH, GAPDH, IDH, PDH, ADH, whilst active oxidative burst was implied when NAD(P)H was bound to NOX enzyme. Based on the similarity of the NAD(P)H fluorescence lifetimes (see Table 1) MDH, HADH, LDH, GAPDH enzymes (1.24 ns – 2.005 ns) were in the following collectively classified as “LDH-like”, associated with anaerobic glycolysis. IDH, PDH, ADH (2.17 ns – 2.60 ns) were grouped into “PDH-like” enzymes, associated with mitochondrial metabolism (especially OxPhos). NADPH bound to NADPH oxidases (NOX) with their long fluorescence lifetime have an own class “NOX”. Further, proportion of the pixel area of the assigned enzyme group in the pixel area of the segmented cell (fraction/cell area) was calculated^69^.

#### 2. *Ascaris suum* larval segmentation

At first glance, the images of *A. suum* larvae showed two regions with different NAD(P)H FLIM properties. The NAD(P)H fluorescence intensity of the larvae within the posterior region was brighter compared to that of the anterior region (Supplementary Fig. S1a, first column). The fluorescence lifetime images with false color representation indicated varying NAD(P)H lifetimes within the larvae (Supplementary Fig. S1a, second column). The anterior region that was inclusive of the internal and external section of the larvae, were referred to as pharynx, and posterior region more internal to the body, were referred to as midgut ^70^ (Supplementary Fig. S1a, third column). Given the numerical aperture of the microscope objective lens, anatomical details of these compartments in the larvae could not be resolved. Lifetimes were allocated to binding of the co-enzyme to its respective partners in the phase domain, with enzyme vectors pointing from unbound “free” NAD(P)H to enzyme-bound NAD(P)H, as depicted by the phasor plot (Supplementary Fig. S1b). The allocated enzymes and enzyme vector lengths were assigned respective colors (color key scale with respective enzyme names and percentage activity on the right of Supplementary Fig. S1b) allowing for the visualization of the enzyme and activity map, respectively, within the larval regions.

In order to analyze the two regions systematically, we separated the *A. suum* images in these two regions by performing image segmentation using the fast random forest-based pixel classification algorithm LABKIT (Fiji plugin, https://imagej.net/plugins/labkit/ downloaded 16th March 2023) ^71^. The algorithm was trained by manually identifying the pharynx, midgut, and background regions in some of the NAD(P)H intensity images, afterwards the trained algorithm was applied to the remaining images. In each FoV, the segmentation resulted in two binary masks for the two annotated image regions. We multiplied both the enzyme and activity maps of each FoV by the corresponding binary masks for the regions 1 and 2 (Supplementary Fig. S1c and S1d). The activity gives the ratio of enzyme-bound NAD(P)H to total NAD(P)H: 0% indicating metabolic inactivity (only free NAD(P)H) to 100% indicating highly active, complete binding to the enzymes. The percentual enzyme-bound NAD(P)H activity is represented in maps (Supplementary Fig. S1d).

The segmented activity maps were averaged over the entire respected body region, resulting in a mean free: enzyme-bound NAD(P)H “activity” value for every larva for each anterior and posterior region; at least ten larvae were imaged per treatment. To analyze the enzyme maps their histograms were generated for anterior and posterior region of the larvae and normalized for the pixel area resulting in fraction of the respected enzyme per area. Similar to the macrophages, enzymes with similar NADH(P)H fluorescence lifetimes were grouped and proportion of the pixel area of the assigned enzyme group in the pixel area of the segmented body region(fraction/body region) was calculated.

### Statistical data analysis

Statistical analysis and visualization were done in R (version 4.4.0) ^72^ and Prism (version 5, GraphPad Software, Inc.). The R package Seurat ^73^ was adapted for the UMAP analysis of the single cell NAD(P)H-FLIM macrophage data. Macrophage data were transformed by dividing each value by the total value of the cell and multiplied by the default scale factor of 10^4^. Thereafter, natural log with a pseudo-count of 1 was applied. Dimension reduction was done using principal component analysis (PCA) and clusters identified by a shared nearest neighbour modularity optimization clustering algorithm. Uniform manifold approximation and projection (UMAP) via the R package uwot ^74^ was used for dimension reduction and spatial visualization. Receiver operating characteristic (ROC) analysis was used for cluster and condition comparisons. P value was adjusted for multiple comparison using the Bonferroni method. A log fold change of 1 with a p. adjusted value of <0.001 was applied to distinguish effects of cytokine polarisation or drug on the metabolic profiles of the macrophages.

For larval data, baseline activity was determined by comparing the two regions of larvae unexposed to ARTs or DMSO using the Wilcoxon statistical and effect size tests. Unsupervised PCA with k-means clustering was done for dimension reduction and cluster identification. Further, supervised UMAP using the approximate nearest neighbor (‘annoy’) method was done for spatial visualization. To determine impact of ART exposure on activity across all groups, Kruskal wallis was used. As all drugs were dissolved in DMSO, statistical analysis was done in comparison to the DMSO vehicle control.

## Results

### High OxPhos/low anaerobic glycolysis metabolic profile of resting porcine alveolar-derived macrophage cell line

Macrophages can alter their metabolic profiles in response to cytokine stimulation or the surrounding metabolic environment. When resting macrophages (M0) are stimulated with proinflammatory cytokines such as IFN-γ, they polarize towards pro-inflammatory (M1-like) macrophages marked by a metabolic profile with elevated anaerobic glycolysis, a disrupted TCA cycle, and the expression of inducible nitric oxide synthase ^75–77^. Conversely, stimulation with anti-inflammatory cytokines like IL-4 causes M0 macrophages to switch to anti-inflammatory (M2-like) macrophages, which exhibit a canonical, OxPhos-based metabolic profile ^75–77^. Macrophage cell lines, due to their proliferation and immortalization, may show altered energy metabolism compared to primary macrophages. Given that macrophages used in this study were derived from the PAM 3D4/31 cell line, NAD(P)H-FLIM was used to investigate the steady-state metabolic profile of the cell line under unstimulated conditions.

The fluorescence intensity of the macrophages in medium (M0) (Fig. 1a, first column) and fluorescence lifetime images with false color representation indicating NAD(P)H lifetimes within the macrophages (Fig. 1a, second column) were determined and analyzed by our phasor approach (Fig. 1a, third column). The allocated enzymes and enzyme vector lengths were assigned respective colors (color key scale with respective enzyme abbreviation and percentage activity at the bottom of Fig. 1a, fourth and fifth column) allowing for the visualization of the enzyme and activity map, respectively.

**Fig. 1.**
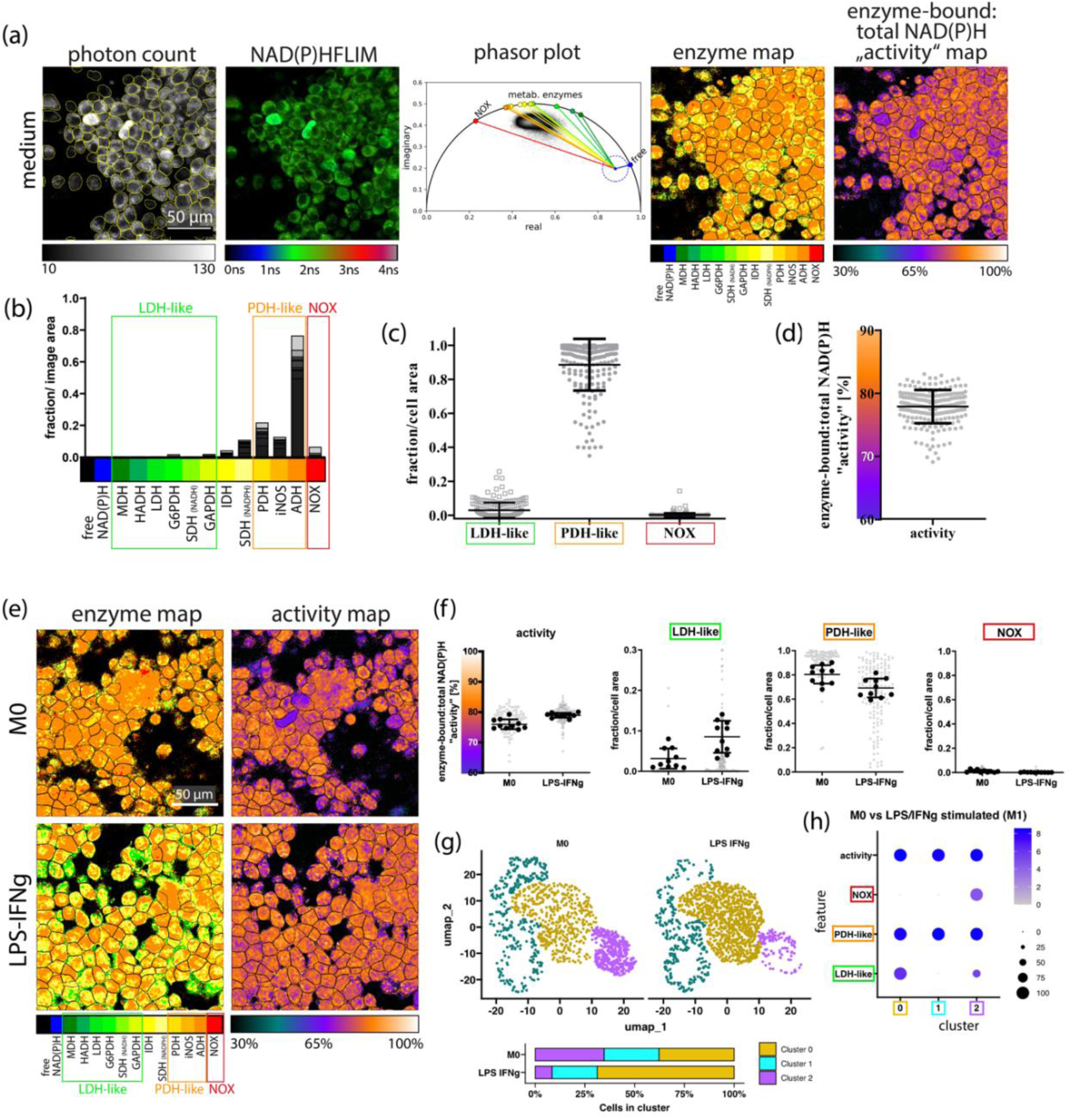
Metabolic profile of M0 and M1-like porcine alveolar-derived macrophage cell line. **(a)** Representative microscopic image (one from ten) of M0 macrophages. From left to right: NAD(P)H fluorescence intensity – the scale bar indicates 50 µm and applies for all shown images; NAD(P)H fluorescence lifetime image in false colour representation ranging from 0 – 4 ns; corresponding phasor plot with enzyme vectors pointing from unbound “free” NAD(P)H to NAD(P)H bound to various enzymes, map of the allocated NAD(P)H-dependent enzymes; map of the enzyme-bound: total NAD(P)H activity in false colour representation ranging from 30 – 100%. **(b)** Superimposed enzyme histograms of ten FoVs, each averaged over the entire FoV with “LDH-like”, “PDH-like” and “NOX” groupings in M0 macrophages. **(c)** Plotted proportion of the pixel area of the assigned enzyme group in the pixel area (fraction/cell area) of the segmented M0 macrophages. **(d)** Averaged enzyme-bound: total NAD(P)H activity of the segmented M0 macrophages (c & d) Each gray dot represents a single cell. **(e)** Representative microscopic image of the enzyme map and activity map of resting M0 and LPS/IFNy polarised (M1-like) macrophages. **(f)** Dot plots depicting overall enzyme activity, LDH-like enzymes, PDH-like enzymes and NOX of M0 and LPS/IFNy. Each gray dot represents a single cells in 1 FoV while the black dots represent average over FoV (5 in 2 wells). **(g)** UMAP plot illustrating clusters 0, 1 and 2 (represented by goldenrod, cyan and purple respectively) within M0 and LPS/IFNy polarised macrophages. The stacked barplot denotes the percentage distribution of cells withing each cluster in M0 and LPS/IFNy. **(h)** The dot plot illustrates the metabolic profiles (features) associated with each cluster. Features on the y axis and the clusters on the x axis. The increasing grey-blue gradient indicates the intensity of the metabolic profile within the cluster while the black dots denote the percentage size of cells associated with the metabolic profile within the cluster.

Based on the similarity of the NAD(P)H fluorescence lifetimes (see Table 1) MDH, HADH, LDH, GAPDH enzymes were grouped into LDH-like enzymes and IDH, PDH, ADH were grouped into PDH-like enzymes denoting anaerobic glycolysis and OxPhos respectively (Fig. 1b). Analysis of the M0 macrophages revealed an energy metabolic profile of high aerobic i.e. OxPhos/low anaerobe glycolysis as depicted by the high composition of PDH-like enzymes and low composition of LDH-like enzymes (Fig. 1c). Also observed was the low composition of enzymes involved in oxidative burst (NOX). At steady state, the M0 macrophages were observed to have an overall metabolic activity of ∼77% (Fig. 1d).

Next we evaluated the metabolic profile of IL-4 stimulated (M2-like) macrophages. The overall metabolic activity of the M2-like macrophages was comparable to that of M0 macrophages. As expected, the energy metabolic profile of M2-like macrophages appeared to be comparable to that of the M0 macrophages (Supplementary Fig. S2a), displaying a metabolic profile of high Oxphos (high PDH-like enyzmes) and low anaerobic glycolysis (low LDH-like enzymes); with a slight increase in PDH-like enzyme activity (Supplementary Fig. S2b). Single-cell cluster analysis classified the M0 vs M2-like macrophages into three clusters (Supplementary Fig. S2c). Cluster 0 depicted by goldenrod, displayed both high PDH-like and high LDH-like enzyme activity, indicative of a combined high OxPhos and high anaerobic glycolysis metabolic profile whilst cluster 1 (darkcyan) demonstrated NOX activity, low LDH-like activity and high PDH-like activity indicative of an oxidative burst metabolic profile and an energy profile of high Oxphos/ low anaerobe glycolysis (Supplementary Fig. S2c and 2d). Cells in cluster 2 (purple) displayed an exclusive inclination towards PDH-like enzyme activity indicative of high OxPhos (Supplementary Fig. S2c and 2d). Markedly, a negligible energy metabolic shift was observed in M2-like macrophages from combined high OxPhos and high anaerobic glycolysis (goldenrod cluster) towards the high OxPhos/ low anaerobic glycolysis (darkcyan) (Supplementary Fig. S2c); an observation complemented by the slight increase in PDH-like enzyme activity (Supplementary Fig. S2b).

Given that the energy metabolic profile of the M0 and M2-like macrophages are rather similar (Supplementary Fig. S2), we explored the ability of NAD(P)H FLIM to identify metabolic shifts triggered by LPS/IFN-γ stimulation in this macrophage cell line. M1 polarisation had minimal influence on the overall metabolic activity of the cell line (79±1)% in comparison to that of M0 macrophages (76±2)% (Fig. 1e and 1f). However, the enery metabolism changed with an increase in anaerobic glycolysis (LDH-like enzymes) and reduction in Oxphos (PDH-like enyzmes) (Fig. 1e and 1f). Single-cell clustering identified three clusters in M0 vs M1-like macrophages (Fig. 1g). Cluster 0 (goldenrod) exhibited both high PDH-like and high LDH-like enzyme activity, indicative of a combined high OxPhos and high anaerobic glycolysis metabolic profile (Fig. 1g and 1h). Exclusively high OxPhos was observed in cluster 1(darkcyan) marked by the high PDH-like enzyme activity and absence of LDH-like enzyme activity (Fig. 1g and 1h). Lastly, cluster 2 (purple) was observed to have a combined high Oxphos/ low anaerobic glycolysis and oxidative burst metabolic profile indicated by the high PDH-like enzyme activity/ low LDH-like enzyme activity and NOX enzyme activity (Fig 1g and 1h). Notably, LPS/IFN-γ induced an accumulation of LDH-like enzyme activity in the macrophages (∼1.38-fold; AUC = 0.632) causing an energy metabolic shift from high OxPhos/ low anaerobe glycolysis state (purple cluster) to a high Oxphos/ high anaerobic glycolysis state (goldenrod cluster) (Fig. 1g).

Taken together, resting (M0) PAM 3D4/31 cells exhibited a metabolic profile of high OxPhos/ low anaerobe glycolysis with an overall metabolic activity of 76%. Consistent with findings in M1-polarized human macrophages ^78^, M0 macrophages polarized to the M1 phenotype with LPS/IFN-γ showed increased LDH-like enzyme activity and decreased PDH-like enzyme activity. This hints to an increased cytosolic metabolism including lactate formation (anaerobic glycolysis), while the mitochondrial metablism is still active but reduced compared to unstimulated macrophages.

### Artesunate enhances LDH-like enzyme activity and reduces overall metabolic activity of M2-like macrophages

*Ascaris* larval migration through the lungs induces a localised anti-inflammatory cytokine profile characterized by IL-4 that polarize macrophages within the lung towards an M2-like phenotype ^9,11,79^. We evaluated the impact of artesunate on M2-like lung macrophages by analyzing the metabolic profiles of IL-4 polarised macrophages after a 24-hour exposure to 1 µM artesunate. Using two-photon NAD(P)H-FLIM imaging, we observed an increase in LDH-like enzymes (indicating anaerobic glycolysis), a decrease in PDH-like enzymes (oxidative mitochondrial metabolism especially OxPhos), and a reduction in overall metabolic activity (73±1)% in artesunate-treated M2 macrophages compared to their respective DMSO controls (77±1)% (Fig. 2a and 2b).

**Fig. 2.**
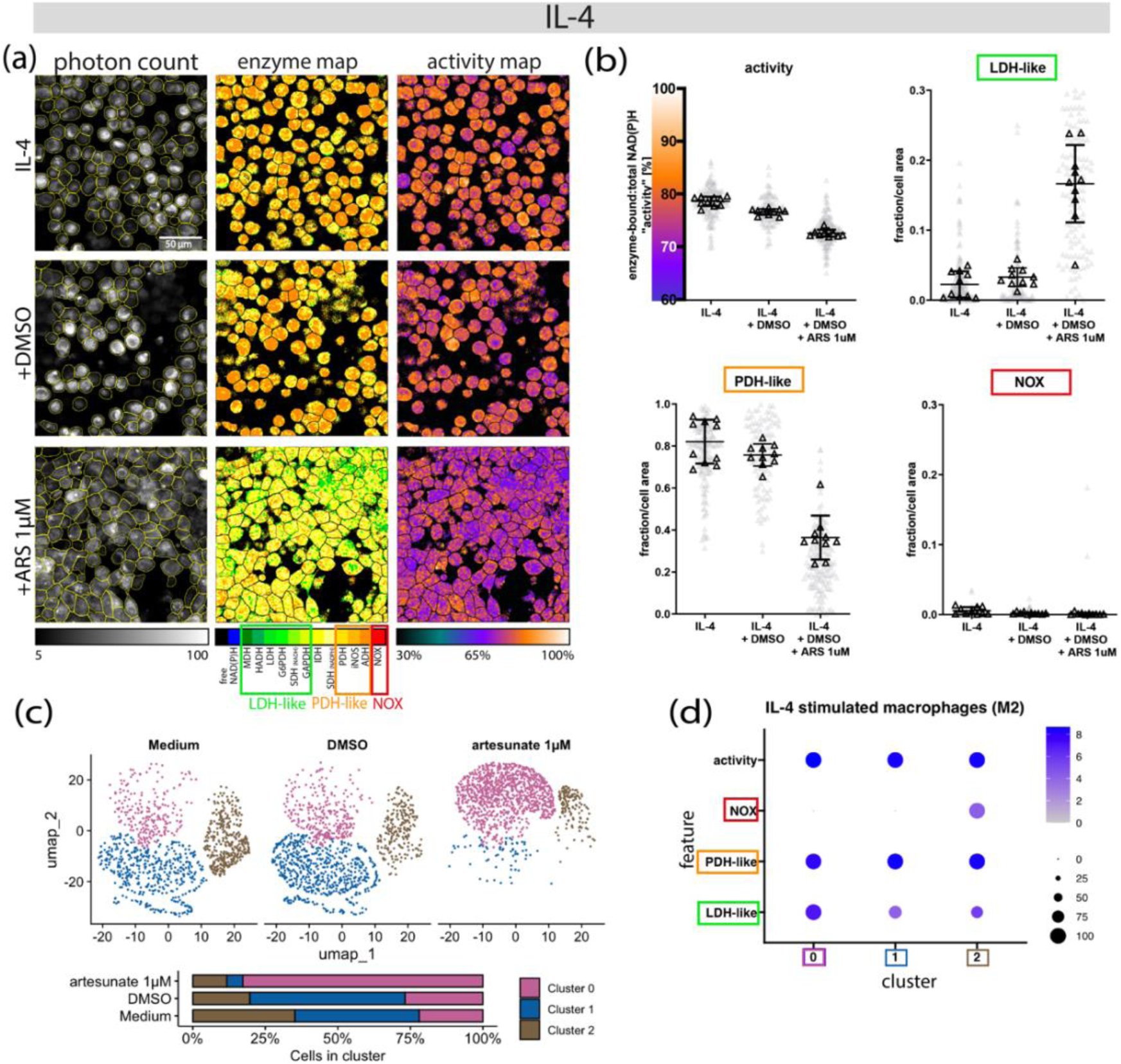
Artesunate increases LDH-like activity and decreases the overall metabolic activity of M2-like macrophages. Porcine M2-like macrophages were exposed to 1 μM of artesunate (ARS) for 24 hours and evaluated using two-photon NAD(P)H-FLIM. **a**) Representative microscopic image of M2-like macrophages. Left: map of the allocated NAD(P)H-dependent enzymes; right: map of the enzyme-bound: total NAD(P)H activity in false colour representation ranging from 30 – 100%. (**b**) Dotplots illustrating the overall metabolic activity, LDH-like, PDH-like and NOX enzyme activity respectively of IL-4 stimulated macrophages exposed to medium, DMSO and artesunate respectively. **(c)** The UMAP plot illustrates clusters of single cells exposed and unexposed to artesunate. Cluster 0, 1 and 2 are respresented by magenta-pink, blue and brown respectively. The stacked barplot denotes the percentage distribution of cells withing each cluster in respective conditions: Medium, DMSO and artesunate 1µM. **(d)** The dot plot illustrates the metabolic profiles (features) associated with each cluster. Features on the y axis and the clusters on the x axis. The increasing grey-blue gradient indicates the intensity of the metabolic profile within the cluster while the black dots denote the percentage size of cells associated with the metabolic profile within the cluster.

Single-cell clustering identified three clusters in the M2-like macrophages, all predominantly associated with high PDH-like enzyme activity (Fig 2c and 2d). Cluster 0 (magenta-pink) exhibited high LDH-like enzyme activity, indicative of a high OxPhos/ high anaerobic glycolysis metabolic profile (Fig 2c and d). In contrast, cluster 1 (blue) displayed low LDH-like enzyme activity, reflecting a high OxPhos/low anaerobic glycolysis metabolic profile (Fig 2c and d). Notably, the blue cluster showed a ∼ 4.24-fold reduction (AUC =0.018) relative to the magenta-pink cluster. Lastly, cluster 2 (brown) demonstrated low LDH-like enzymes and low NOX enzymes (Fig 2c and d).

Parallel to the M2-like macrophages exposed to artesunate, we evaluated the effect of the drugs on M0 macrophages metabolism. Comparably, M0 macrophages exposed to artesunate displayed an increase in LDH-like enzymes, a decrease in PDH-like enzymes, and a reduction in overall metabolic activity (78±1)% to (73±1)%) (Supplementary Fig. S3a and S3b). While four clusters were identified in M0 macrophages, cluster 0 (magenta-pink), 1 (blue) and 2 (brown) (supplementary Fig. 2c) displayed similar metabolic profiles as respective clusters of M2-like macrophages (Fig 2c and 2d). The M0 blue cluster, similarly, exhibited a ∼ 3.7-fold reduction in LDH-like enzymes (AUC = 0.036) relative to the magenta-pink cluster. In contrast to M2-like macrophages, cluster 3 (grey) was exclusive to M0 macrophages and exhibited high PDH-like enzyme activity (Supplementary Fig. S3c and S3d).

M0 and IL-4 stimulated M2-like macrophages predominantly exhibited a high OxPhos/low anaerobic glycolysis metabolic profile, represented by the blue cluster (Fig 2c; Supplementary Fig. S3c). However, artesunate exposure induced an accumulation of LDH-like enzymes in M0 (L2FC = 1.86; AUC = 0.761) and M2-like macrophages (L2FC = 2.41; AUC = 0.867) relative to DMSO control, shifting the metabolic profile towards high OxPhos/high anaerobic glycolysis, as evidenced by an increase in cells within the magenta-pink clusters and a corresponding decrease in the blue clusters (Fig 2c; Supplementary Fig. S3c.). Regardless of culture conditions, NOX enzyme activity (brown cluster) remained low and was observed in only a small subset of macrophages, suggesting minimal oxidative burst activity (Fig 2c; Supplementary Fig. S3c.).

Collectively, exposure to artesunate induced a shift in the predominant profile from high OxPhos/ low anaerobic glycolysis (blue cluster) to high OxPhos/ high anaerobic glycolysis (magenta-pink cluster) through LDH-like enzyme accumulation in both M0 and M2-like macrophages, along with a reduction in overall metabolic activity compared to DMSO controls. NOX enzyme activity (brown cluster) remained minimal, indicating negligible oxidative burst activity across conditions.

### Distinct metabolic profiles within anatomical regions of *A. suum* L3

Next, we applied NAD(P)H-FLIM to analyze the metabolism of *A. suum* L3 larvae and identified two distinct metabolic profiles that were anatomically localized (Fig. 3a). Specifically, we observed a clear difference in both the bound fluorescence lifetime and fluorescence intensity between the pharynx and midgut regions of the larvae (for the anatomy of *Ascaris* L3, see Supplementary Fig. S1a). Analysis of the overall metabolic activity maps of larvae from the control conditions (artesunate unexposed larvae) revealed a significantly higher (p<0.0001) NAD(P)H dependent overall metabolic activity in the pharynx region (72 ± 3%) compared to the midgut region of the larvae (58 ± 5%) (Fig. 3b) with a large effect size r of 0.817. Additionally, the preferential NAD(P)H-dependent enzyme activity differs between these two regions of *A. suum* larvae (Fig. 3c).

**Fig. 3.**
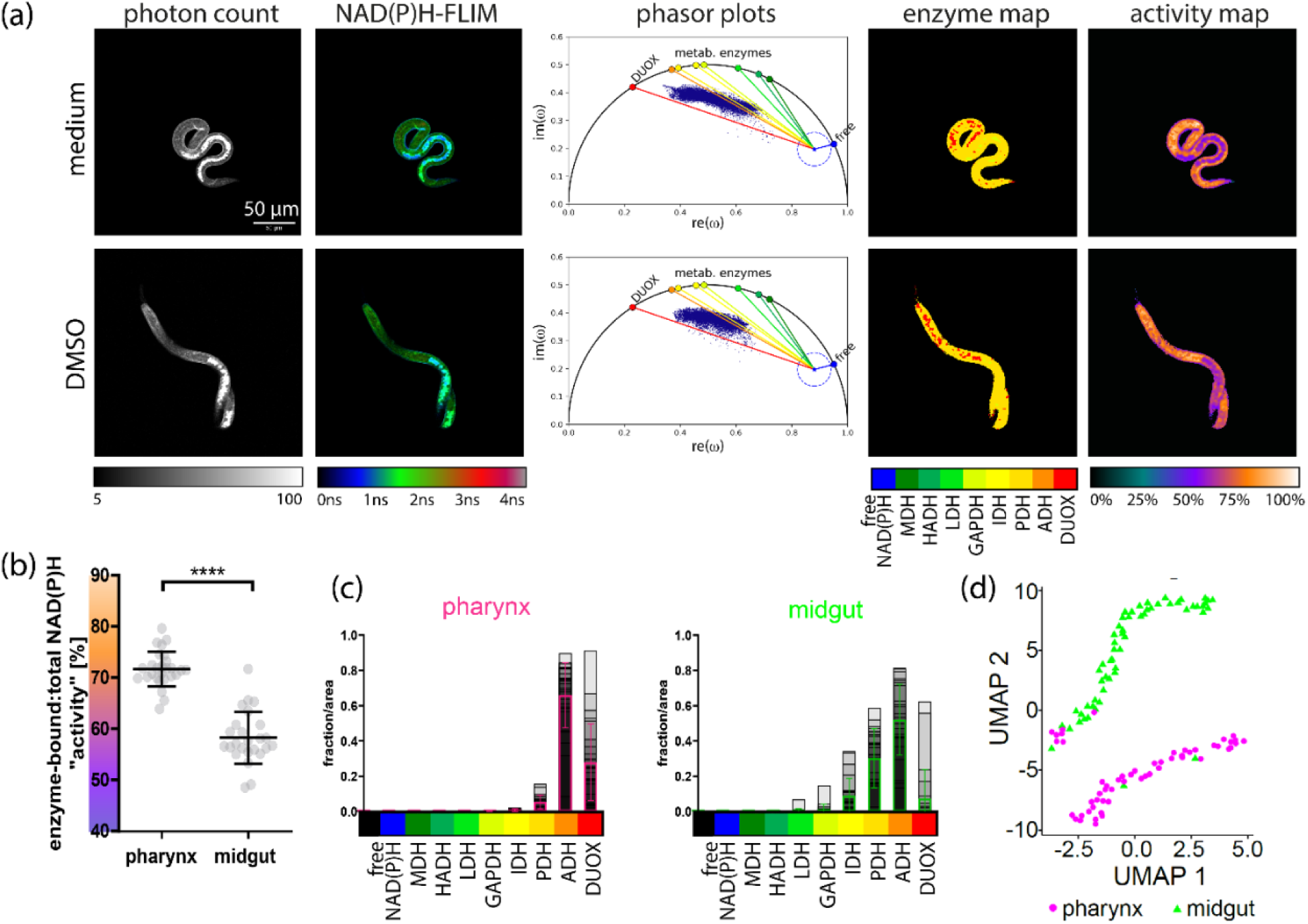
Spatially distinguishable NAD(P)H-dependent metabolic fingerprints in *A. suum* L3. A representative microscopic image of L3 **(a)** visually illustrating the NAD(P)H intensities, NAD(P)H-FLIM, phasor plots, enzyme maps and activity maps in ART unexposed larvae. The scatterplot **(b)** denotes NAD(P)H enzyme activity (%) with standard deviation within the pharynx (circles) and midgut (triangles) regions of larvae unexposed to ARTs. Activity was calculated as the ratio of enzyme bound NAD(P)H to total NAD(P)H; 0% indicating enzyme inactivity (only free NAD(P)H) to 100% indicating highly active, NAD(P)H completely bound to the enzymes). The plots **(c)** denote enzyme histograms normalized to pixel area for pharynx and midgut; grey to black show the overlapped histograms of all larvae, the magenta (pharynx) and green (midgut) bars show the average over all images with standard deviation. UMAP plot **(d)** illustrates the spatial distribution of the allocated bound enzymes and the enzyme-bound: total activity within the pharynx and midgut (represented by magenta dots and green triangles respectively) of ART unexposed larvae. Scale bar indicates 50 µm and applies for all shown images. Significance is represented by p<0.0001: ****; Wilcoxon test.

Despite its lower overall metabolic activity, the midgut exhibited low levels of bound LDH-like and high levels of PDH-like enzymes, here GAPDH, IDH, and PDH, which are associated with bioenergetics. In contrast, the pharynx region appeared to have higher DUOX activity compared to the midgut (Fig. 3c). A K-nearest neighbor (KNN) supervised analysis further highlighted a distinct spatial separation between the pharynx and midgut, consistent with the observed differences in enzyme activity distribution and overall metabolic activity (Fig. 3d).

### Unperturbed metabolic profiles of pharynx and midgut regions of *A. suum* L3 upon exposure to ARTs

The viability of the ART-exposed larvae was investigated using the resazurin reduction assay and ATP bioluminescence assay. The capacity of the resazurin reduction assay to detect the metabolic activity of *Ascaris* larvae was assessed previously^54^. Only high concentrations of ARTs decreased the viability of the larvae after 48 hours of exposure. Due to the more potent effects of artemether on the general metabolic activity (Supplementary Fig. S4) and artesunate on ATP levels (Supplementary Fig. S5), investigations in FLIM focused on exposures of the larvae to 20- and 200 µM artesunate and artemether only. Exposures were done for both 24- and 48-hour timepoints.

The NAD(P)H-dependent metabolic profile and overall metabolic activity of ART-exposed larvae were assessed using the two-photon NAD(P)H-FLIM approach (Fig. 4a). We found the overall metabolic activity of artesunate- and artemether-exposed larvae to be comparable to that of the controls, after 24 hours (Fig. 4b). By applying the silhouette method, three optimal clusters were identified (Supplementary Fig. S6a) and subsequently utilized in K-means unsupervised clustering to categorize the larvae into metabolically distinct groups. These clusters were then correlated with specific metabolic profiles. Cluster 1 (magenta-pink) was associated with oxidative burst characterized by a DUOX activity, exhibited by some larvae (Fig. 4c; Supplementary Fig. S6b). Energy production of the larvae was associated with clusters 2 (brown) and 3 (blue) marked by LDH-like and PDH-like enzymes respectively (Fig. 4c). Only a few larvae exhibited LDH-like enzyme activity (Supplementary Fig. S6c) with majority demonstrating PDH-like enzyme activity (Supplementary Fig. S6d) further highlighting a preference towards OxPhos for bioenergetics.

**Fig. 4.**
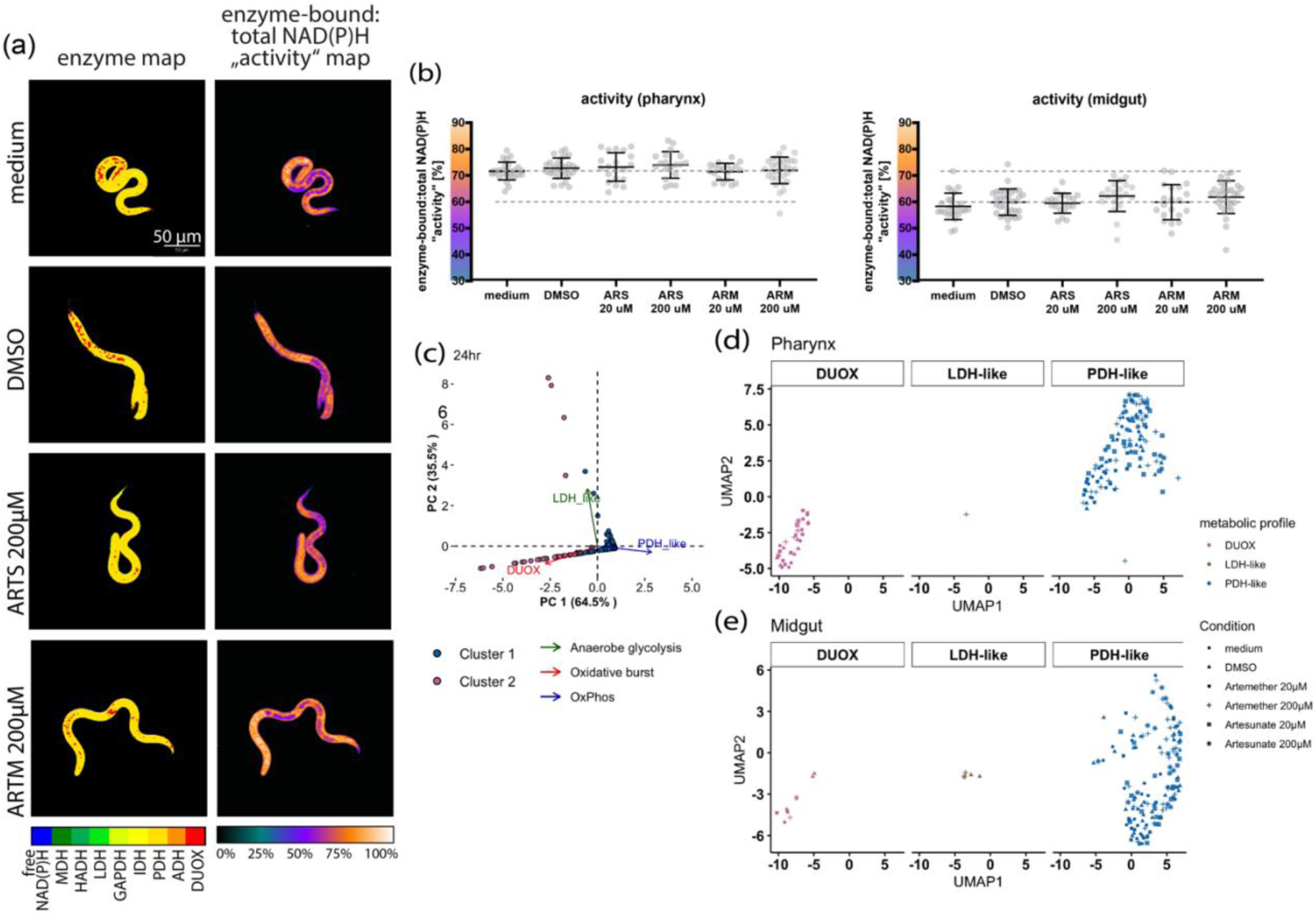
Un-perturbed NAD(P)H-dependent metabolism in ARTs-exposed *A. suum* L3. *A. suum* L3 were exposed to 200 μM of artesunate (ARTS) and artemether (ARTM) for 24 hours and evaluated using two-photon NAD(P)H-FLIM. A representative microscopic image of L3 **(a)** visually illustrating the NAD(P)H enzyme map and activity map in ART exposed larvae. The dot plots **(b)** depicting enzyme activity within the pharynx and midgut. Activity is calculated as the ratio of enzyme bound NAD(P)H to total NAD(P)H; 0% indicating enzyme inactivity (only free NAD(P)H) to 100% indicating highly active, NAD(P)H completely bound to the enzymes). Data dimension reduction was done using PCA and transformed data used for downstream analysis. The PCA biplot **(c)** represents the association between the clusters and the bound enzymes within PC1 and PC2. Cluster 1, 2 and 3 are represented by magenta-pink, brown and blue dots respectively whilst enzymes associated with LDH-like, PDH_like and DUOX-like enzyme activity are depicted in green, blue and red arrows and texts respectively. The UMAP plot and barplots illustrating distribution of metabolic profiles within the pharynx **(d)** and midgut **(e)** of the larvae. Larvae exposed to artemether and artesunate at concentrations of 200- and 20 µM, DMSO and medium are represented by a star, an x marked square, a plus sign, a square, a triangle and a dot respectively. n Artesunate 200/20 µM = 38/40; n Artemether 200/20 µM = 58/40; n medium/DMSO controls = 52/60.

The metabolic profiles specific to anatomical regions were further analyzed spatially using the UMAP method. While both the midgut and pharynx displayed high PDH-like/low LDH-like enzyme activity indicative of energy production, only the pharynx displayed a greater inclination toward oxidative burst, characterized by the DUOX-like metabolic profile (Fig. 4d and 4e). Despite the metabolic differences between these regions, neither artesunate nor artemether induced a shift in the metabolic profile of the larvae. This is evident from the heterogeneous distribution of both ART-exposed and unexposed larvae across the LDH-like, PDH-like and DUOX-like enzyme activities in the pharynx (Fig. 4d) and midgut (Fig. 4e) regions. However, artesunate and artemether appeared to slightly reduce DUOX enzyme activity in the pharynx in comparison to the untreated controls (Supplementary Fig. S6e).

After 48 hours, DUOX enzyme activity increased in the larval midguts across all conditions, as shown by red spots on the enzyme map (Supplementary Fig. S7a). Overall metabolic activity remained comparable between exposed and unexposed larvae, however, metabolic activity in the larval midgut showed a decreasing trend after 48 hours of ART exposure compared to the increase observed after 24 hours of ART exposure (Supplementary Fig. S7b). Two key clusters were identified: bioenergetics (cluster 1, blue) and oxidative burst (cluster 2, magenta-pink), with bioenergetics dominated by PDH-like over LDH-like enzyme activity (Supplementary Fig. S7c). Spatial analysis revealed DUOX activity in the midgut, which increased with artemether exposure but decreased with artesunate (Supplementary Fig. S7d).

Taken together, neither artesunate nor artemether impacted the metabolic profile and overall metabolic activity of the larvae in respective anatomical regions after 24 hours of exposure. While oxidative burst was primarily associated with the pharynx at the 24-hour timepoint, the midgut similarly displayed oxidative burst along side the pharynx at the 48-hour timepoint. Although not significant, larvae exposed to artemether for 48 hours showed a tendency of an increase in DUOX-like enzyme activity in the pharynx.

As summarized in Fig. 5, the midgut section of *A. suum* larvae was observed to have a higher activity of NAD(P)H-bound mitochondrial enzymes associated with energy metabolism, indicating a high OxPhos (high PDH-like enyzmes) and low anaerobe glycolysis (low LDH-like enzymes) metabolic profile. Markedly, the pharynx region of the larvae had a higher contribution of DUOX-like enzyme in comparison to the midgut. Porcine macrophages polarized towards an M2 macrophage phenotype were also observed to have a high OxPhos/low anaerobe glycolysis energy metabolism at steady state (Fig. 5). Although ARTs are observed to have an influence on the NAD(P)H-dependent metabolism of host macrophages, exposure to ARTs had no impact on the NAD(P)H-dependent metabolism of the *Ascaris* larvae.

**Fig. 5.**
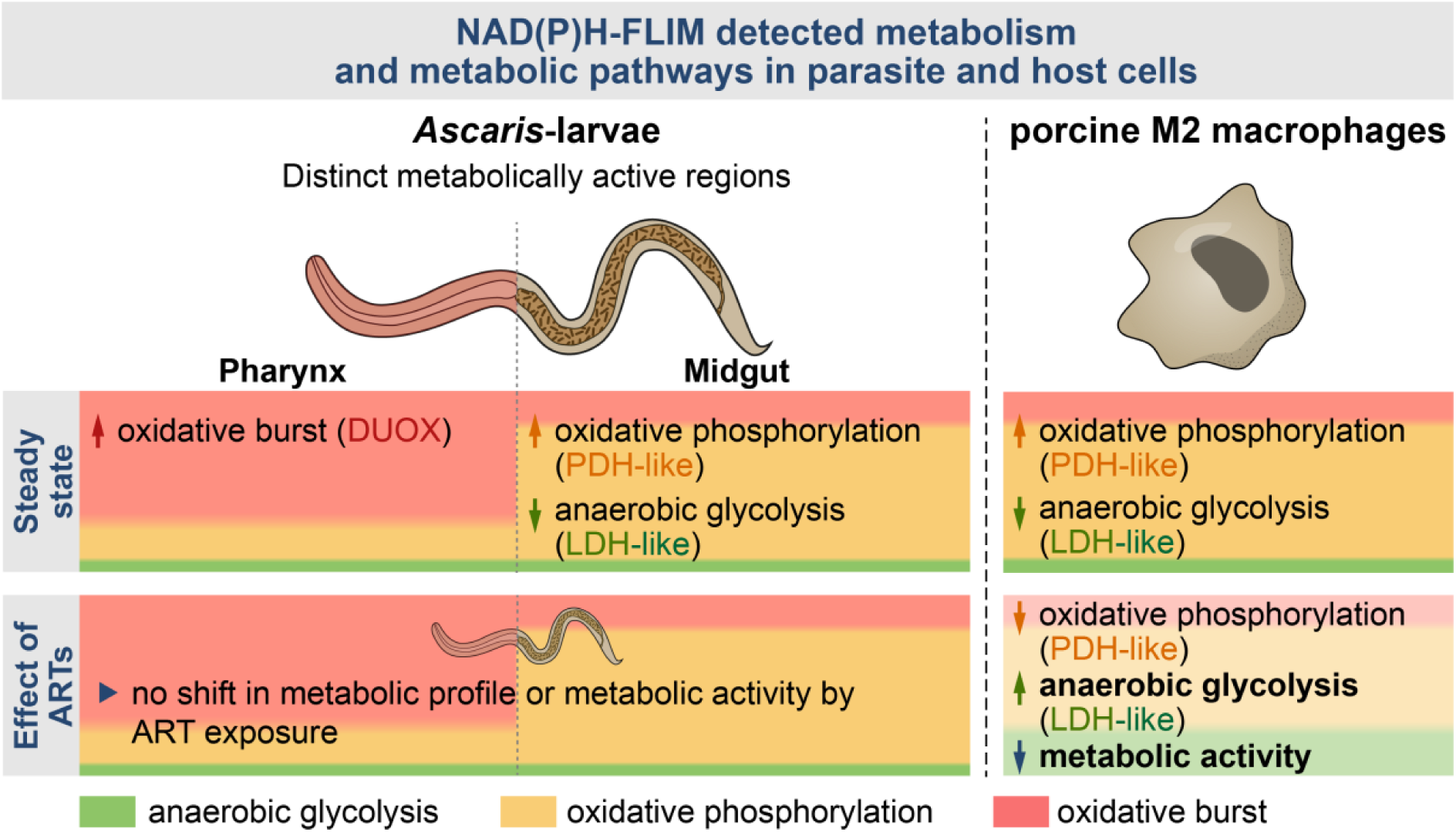
Steady state energy metabolism of high OxPhos / low anaerobe glycolysis in both *Ascaris* larvae and porcine M2 macrophages. NAD(P)H FLIM detected distinct metabolically active regions in the *A. suum* L3. Although both regions were seen to have steady state energy metabolic profiles of high OxPhos (orange) / low anaerobe glycolysis (green), the anterior region referred to as pharynx was highly associated with oxidative burst (pink) when compared to the posterior region referred to as midgut. Similarly, the porcine M2 macrophages had a steady state energy metabolic profile of high OxPhos / low anaerobe glycolysis. Oxidative burst via NOX was however non-existent. Unlike *A. suum* L3, exposure to ARTs decreased the metabolic activity of porcine M2 macrophages and further induced a metabolic shift towards higher anaerobic glycolysis and lower oxidative mitochondrial metabolism.

## Discussion

Co-infections of soil-transmitted helminths (STH) and *Plasmodium falciparum* are common in regions where both parasites are endemic ^15–17^. While the anthelmintic effects of ARTs have been primarily documented for trematodes and cestodes, there has been less emphasis on gastrointestinal nematodes that infect mammals ^80^. Bearing in mind that *A. lumbricoides* is highly prevalent in malaria endemic areas, the potential of ARTs to directly (larval parasite) or indirectly (via host cells) affect ascariasis are conceivable. Given that ARTs are known to target energy metabolism pathways ^31,34,36,81,82^, this study focused on examining the impact of ART on the energy metabolism of *A. suum* larvae and alveolar-derived macrophages. Over the years, fluorescence lifetime imaging (FLIM) has been widely used to study the metabolism of live cells and tissues under drug exposure, providing insights into drug delivery, mechanisms of action, and cellular responses ^47^. NADH and NADPH, collectively termed NAD(P)H due to their overlapping fluorescence spectra, are key cofactors in metabolic reactions, making NAD(P)H-FLIM a powerful tool for real-time analysis of metabolic states at the subcellular level ^45,46^. In this study, two-photon NAD(P)H-FLIM was employed to assess the metabolic profiles of porcine macrophages and *A. suum* L3 larvae, as well as to investigate ART-induced metabolic changes in both host cells and parasites.

Cell lines are developed to allow for rapid proliferation and immortalization of primary cells thereby potentially interfering with their energy metabolism. The porcine macrophage cell line PAM 3D4/31 is reported to rely on both glycolysis and OxPhos ^83^. Consistent with primary alveolar macrophages ^84^, we found that this porcine macrophage cell line exhibits a high OxPhos/low anaerobic glycolysis metabolic profile. This is characterized by elevated activity of mitochondrial PDH-like enzymes and reduced contribution from cytosolic LDH-like enzymes. Macrophage plasticity in immune responses is intricately connected to energy metabolism^77^. Their capacity to modify metabolic profiles in response to the tissue microenvironment, cytokines such as IL-4 or IFN-γ, or antimicrobial agents like LPS is well documented ^75–77,84,85^. The pro-inflammatory M1 phenotype primarily relies on anaerobic glycolysis to meet the immediate energy demands required for inflammatory activity ^75–77^. In line, we found an accumulation of LDH-like enzyme activity in the LPS/IFN-γ-polarised macrophages. In contrast to the M1 phenotype, the anti-inflammatory M2 phenotype is associated with wound healing and infection resolution, and favors OxPhos due to its lower demand for rapid energy production ^75–77^. As expected, polarisation of the M0 macrophages towards an M2-like phenotype elicited no change in the already dominant high OxPhos metabolic profile. At steady-state, most cells utilize OxPhos for their low energy demands ^86^. While macrophages may display comparable metabolic profiles at M0- and M2-like states, the immunological phenotypes, differ with M2-like macrophages characterized by high expression of anti-inflammatory markers such as CD203a, MHCII and arginase 1 ^79,87,88^.

Mitochondria have been implicated in activating ARTs and inducing its mechanism of action ^28,29^. Studies further suggest that ARTs may disrupt electron transport chain activity ^27,30^, potentially causing ATP depletion and mitochondrial dysfunction ^28,29^. Human monocytes polarized to the M2 phenotype have reportedly shifted to an M1 phenotype following treatment with artesunate ^22^. While the study focused on cytokine production and attributed this switch to JAK2/STAT3 pathway inhibition, the findings indicate a potential secondary effect involving metabolic rewiring. In the current study, exposure of the porcine alveolar macrophage cell line to artesunate resulted in accumulation of anaerobic glycolysis with a slight decrease in PDH-like enzymes and overall metabolic activity. Given that a similar effect on the metabolic profile and metabolic activity was demonstrated in M0 macrophages; resting macrophages with an inactive JAK2/STAT3 pathway, we inferred an influence of the drug on the mitochondria of the macrophages.

NAD(P)H-FLIM is a powerful tool for analyzing cellular metabolism across different species, offering valuable insights into parasitism. Using two-photon NAD(P)H-FLIM, we identified two metabolically distinct regions in the larvae: the “pharynx region” and the “midgut region”. Although both regions exhibited OxPhos as the preferential energy metabolic profile, the pharynx displayed a higher level of oxidative burst, associated with a DUOX-like metabolic profile, compared to the midgut. Additionally, our data revealed that the pharynx had higher overall metabolic activity than the midgut. In mammalian cells, enzymes of the NOX family which DUOX is part of, are mainly associated with defense metabolism via oxidative burst in phagocytic cells ^89^, but they have also been linked to non-phagocytic neuronal cells^90^. Given that the pharynx region of *A. suum* L3 (egg-stage) larvae is extensively filled with neuronal cell bodies ^70^, our findings may indicate a highly active nervous system.

In *C. elegans*, DUOX-mediated defense metabolism against bacterial pathogens has been reported in the hypodermis and intestine ^91^, while in *Heligmosomoides polygyrus*, it is mainly observed in the digestive system ^49^. Therefore, the DUOX-like metabolic profile observed in our study, although predominanrtly in the pharynx, suggests defense-related metabolism in *A. suum* larvae. Remarkably, DUOX activity has also been shown to facilitate cross-linking of tyrosine residues between collagens, which is essential for proper biogenesis and function of the cuticle in *C. elegans* ^92^. Since the pharynx is an ectodermal tissue covered with cuticle, while the midgut is mesodermal and lacks a cuticle, it is plausible to hypothesize that the higher DUOX activity in the pharynx may contribute to maintaining cuticle function.

The *A. suum* L3 (egg-stage) midgut consists of seven cells that contain deposits of glycogen and lipids ^70,93,94^ suggesting energy production via glycogenolysis or lipolysis, respectively. However, our analysis revealed that mitochondrial energy metabolism in the midgut is driven by active PDH-like enzymes. The absence of bound HADH indicated that energy production is occurring via aerobic glycolysis rather than fatty acid beta-oxidation, which is typical of lipolysis. Our findings further showed that the larvae’s overall energy metabolism primarily relies on aerobic pathways, marked by the predominant activity of PDH-like mitochondrial enzymes and minimal activity of LDH-like cytosolic enzymes (associated with anaerobic glycolysis). This aligns with the requirement for aerobic respiration during larval development and hepato-tracheal migration^95,96^. This metabolic profile was however not impacted after 24- or 48-hour exposure to ARTs.

Lastly, NAD(P)H-FLIM did not detect any changes in the overall metabolic activity of the larvae, whereas both the resazurin reduction assay and ATP assay (viability assays) demonstrated an ART-induced reduction in larval viability. Notably, the viability assays and the NAD(P)H-FLIM technique are complementary but distinct in their measurements. The resazurin reduction assay measure the NAD(P)H levels of larvae by assessing the oxidation-reduction potential between NAD(P)+ and NAD(P)H ^97^. In contrast, NAD(P)H-FLIM evaluates the activity of NAD(P)H-dependent metabolic catalysis, detecting only the reduced forms NAD(P)H, either free or bound to enzymes. Despite both involving NAD(P)H, the measurements differ fundamentally and are not directly comparable. Similarly, the ATP assay measures ATP levels generated from bioenergetic pathways, while NAD(P)H-FLIM assesses metabolic activity across both bioenergetic and non-bioenergetic pathways ^98^. Although larvae exposed to artesunate exhibited reduced ATP levels, their mitochondria remained functional, as evidenced by active PDH-like enzymes. This indicates that, while ART reduced larval viability after 48 hours of exposure, it did not impact the overall NAD(P)H-dependent metabolic activity associated with both bioenergetic and non-bioenergetic (DUOX-dependent defense) pathways. The higher levels of DUOX activity detected by NAD(P)H-FLIM in the larval midgut after 48 hours as compared to 24 hours suggest increased stress in larvae at this time-point, possibly explaining the low viability measured by the ATP and resazurin reduction assays due to an increased susceptibility to ARTs at this time point.

Collectively, by adapting and applying two-photon NAD(P)H-FLIM to analyze overall metabolic activity and preferential enzymatic activities in porcine macrophages and in *A. suum* L3, we found dominantly mitochondrial energy metabolism in both host and parasitic larvae, characterized by high OxPhos/low anaerobe glycolysis. Surprisingly, we found a high DUOX-like activity in the pharynx of the *A. suum* L3, which we attributed to a possible oxidative burst. Further functions related to DUOX-like activity in *A. suum* larvae need further investigation in future studies. Exposure to artesunate induced a metabolic shift towards higher anaerobic and a reduction in overall metabolic activity in M2-like macrophages. In the context of active larval *Ascaris* infection, we can speculate that a switch from the M2 (high OxPhos/ low anaerobe glycolysis) to M1 (low OxPhos/ high anaerobe glycolysis) macrophage phenotype would delay tissue repair, prolong infection resolution, and hinder larval clearance in the lung, ultimately benefiting the parasite at the host’s expense. Hence, it can be speculated that ART treatments might have an indirect effect on the helminth infection. Our study further found ARTs not to directly affect the metabolic shift in the larval bioenergetics.

The use of two-photon NAD(P)H-FLIM allows for resolution of changes in metabolic profiles and overall metabolic activity, we expect that the method retains the potential to improve our understanding of host-parasite metabolic crosstalk and, by that, of host-parasite interactions in general.

## Acknowledgments

The authors would like to thank Larissa Oser, Philipp Höfler, Beate Anders, Yvonne Weber, Bettina Sonnenburg, Marion Müller, Ralf Uecker, and Christiane Palissa for methodological and technical support. We also thank Anne Winkler for illustrative support.

## Author Contributions

Conceptualization: SH, PHS, SR, ZDM; Investigation: ZDM, RL, AK ; Data curation: ZDM, RL; Formal analysis and Visualization: ZDM, RL; Data interpretation: ZDM, RL, JK, RN, SH; Supervision: SH, RN; Funding acquisition: SH, RN; Resources: SH, RN, AEH; Writing -original draft: ZDM, SH, RL; Writing – review & editing: ZDM, RL, AK, JK, AEH, SR, PHS, RN, SH. All authors have read and agreed to the published version of the manuscript.

## Data availability

All data that support the findings of this study are included within this paper and its FLIM image files are available from the corresponding author upon request.

