## Supplementary for "Two-Photon NAD(P)H-FLIM reveals unperturbed energy metabolism of *Ascaris suum* larvae, in contrast to host macrophages upon artemisinin derivatives exposure"

**Supplementary Material**

**Supplementary Fig. S1**

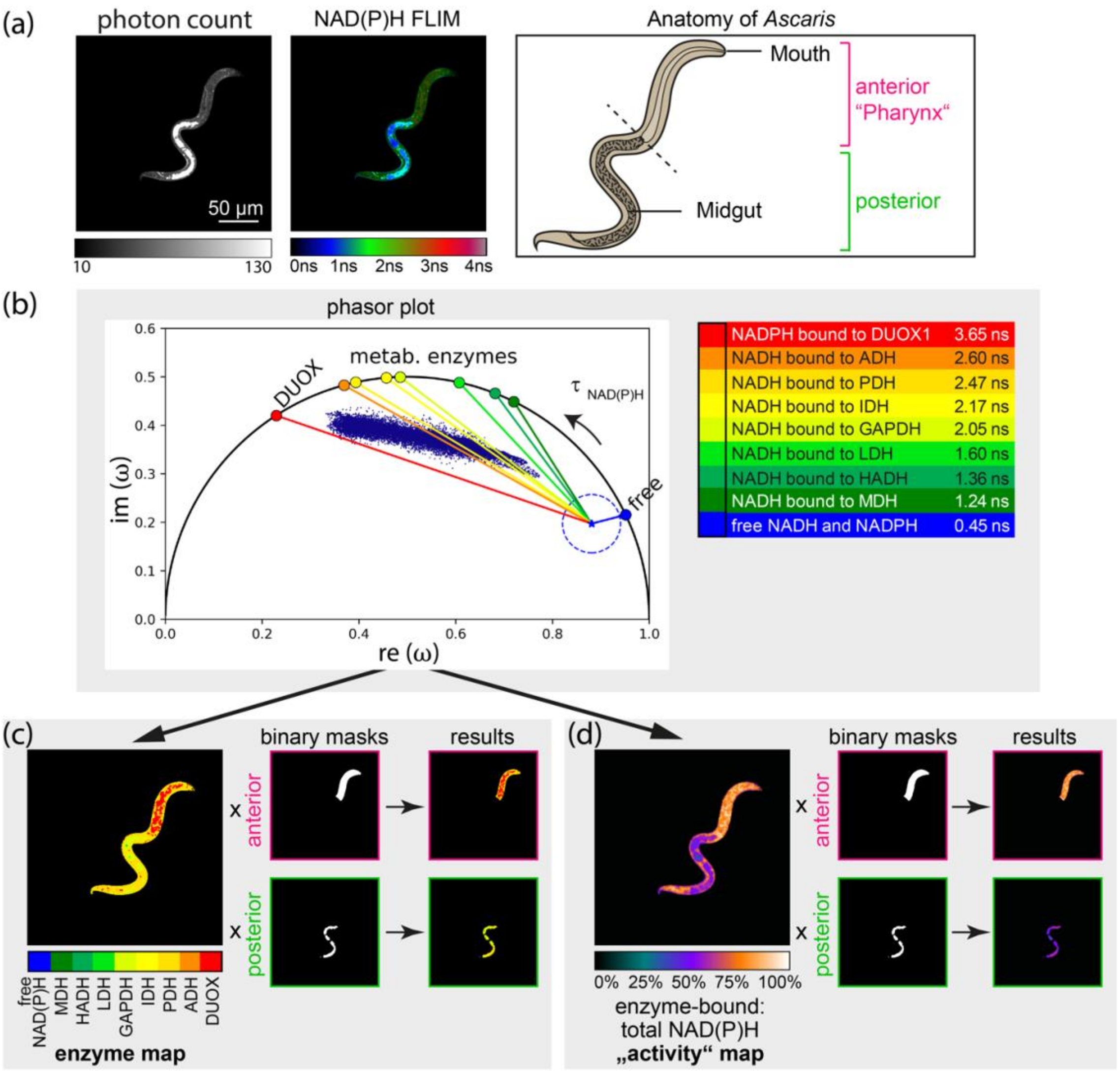


**Fig.S1 Phasor analysis and segmentation of *A. suum* larvae (a)** Form left to right: NAD(P)H fluorescence intensity – the scale bar indicates 50 µm and applies for all shown images, NAD(P)H fluorescence lifetime image in false colour representation ranging from 0 – 4 ns. The anatomy of *Ascaris suum* L3 showing the identified regions. **(b)** Corresponding phasor plot with enzyme vectors pointing from unbound "free" NAD(P)H to NAD(P)H bound to various enzymes. Left: Reference system in phase domain after fast Fourier transformation of NAD(P)H fluorescence lifetimes of free and enzyme-bound NAD(P)H state. Right: List of considered NAD(P)H-dependent enzymes with their specific NAD(P)H-fluorescence lifetimes. (c and d) Workflow of phasor based FLIM analysis [8] and image segmentation by LABKIT, an ImageJ plugin. **(c)** Image segmentation of the enzyme map. **(d)** Image segmentation of the activity map. For both (c) and (d) from left to right: exemplary enzyme (c) or activity (d) map as result of the vector-analysis of the NAD(P)H-FLIM data; corresponding binary masks for anterior (pharynx) and posterior (midgut) body region of the larvae after images segmentation; resulting images per region.

**Supplementary Fig. S2**

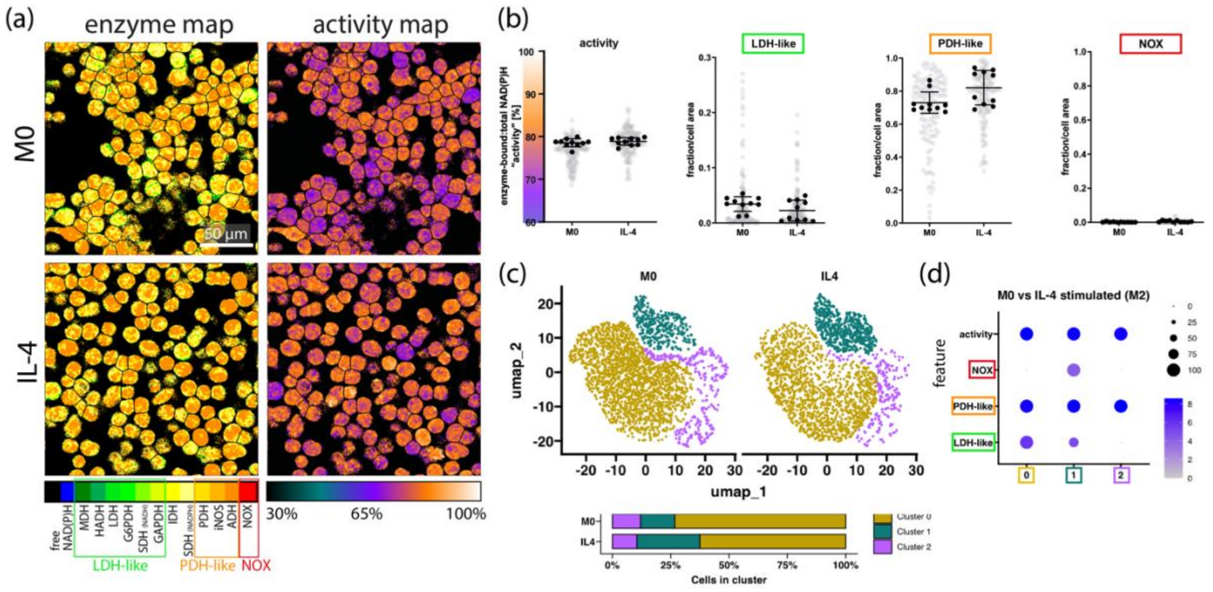


**Fig. S2. Metabolic profile of M0 and M2-like porcine alveolar-derived macrophage cell line (a)** Representative microscopic image of the enzyme map and activity map of resting M0 and IL-4 polarised (M2-like) macrophages. **(b)** Dot plots depicting overall enzyme activity, LDH-like enzymes, PDH-like enzymes and NOX of M0 and IL-4. Each gray dot represents a single cell in 1 FoV while the black dots represent average over FoV (5 in 2 wells). **(c)** UMAP plot illustrating clusters 0, 1 and 2 (represented by goldenrod, dark cyan and purple respectively) within M0 and IL-4 polarised macrophages. The stacked bar plot denotes the percentage distribution of cells withing each cluster in M0 and IL-4. **(d)** The dot plot illustrates the metabolic profiles (features) associated with each cluster. Features on the y axis and the clusters on the x axis. The increasing grey-blue gradient indicates the intensity of the metabolic profile within the cluster while the black dots denote the percentage size of cells associated with the metabolic profile within the cluster.

**Supplementary Fig. S3**

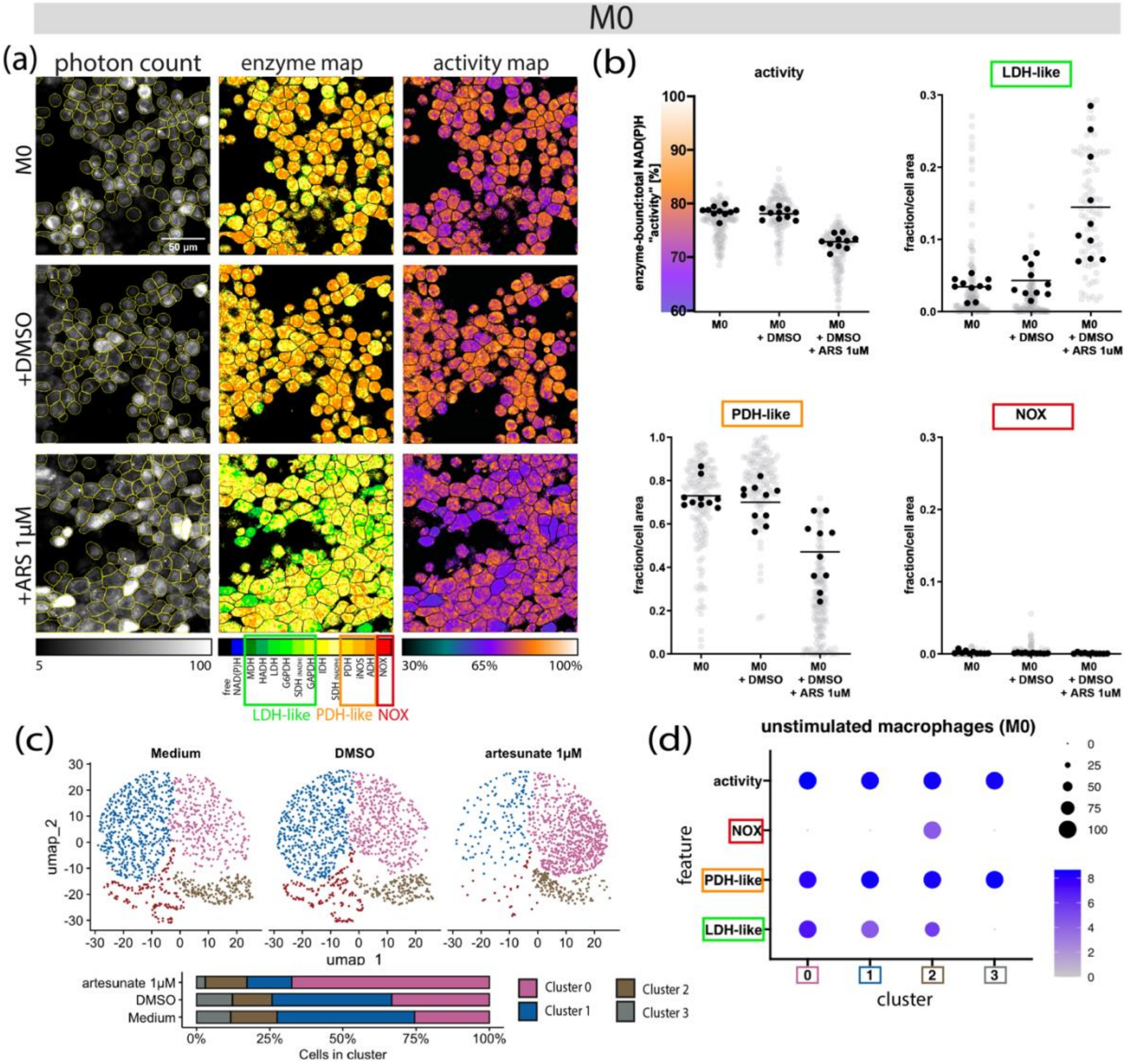


**Fig. S3. Artesunate increases LDH-like activity and decreases the overall metabolic activity of M0 macrophages.** Porcine M0 macrophages were exposed to 1 µM of artesunate (ARS) for 24 hours and evaluated using two-photon NAD(P)H-FLIM. **a**) Representative microscopic image of macrophages. Left: map of the allocated NAD(P)H-dependent enzymes; right: map of the enzyme-bound: total NAD(P)H activity in false colour representation ranging from 30 – 100%. (**b**) Dot plots illustrating the overall metabolic activity, LDH-like, PDH-like and NOX enzyme activity respectively of M0 macrophages exposed to medium, DMSO and artesunate respectively. **(c)** The UMAP plot illustrates clusters of single cells exposed and unexposed to artesunate. Cluster 0, 1, 2 and 3 are represented by magenta-pink, blue, brown and grey respectively. The stacked bar plot denotes the percentage distribution of cells withing each cluster in respective conditions: Medium, DMSO and artesunate 1µM. **(d)** The dot plot illustrates the metabolic profiles (features) associated with each cluster. Features on the y axis and the clusters on the x axis. The increasing grey-blue gradient indicates the intensity of the metabolic profile within the cluster while the black dots denote the percentage size of cells associated with the metabolic profile within the cluster.

**Supplementary Fig. S4**

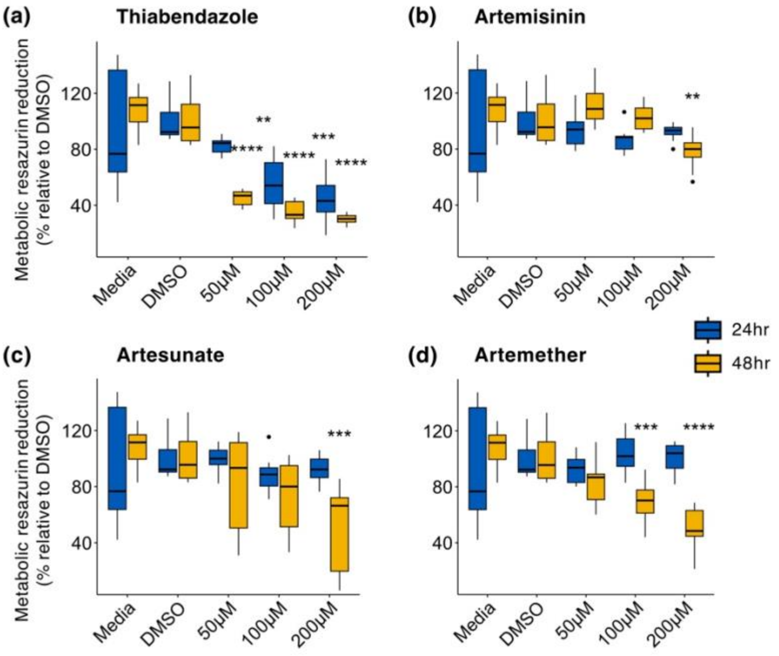


**Fig. S4. 48-hour exposure to artemisinin, artesunate and artemether decreased viability of *A. suum* L3**. In 96 well plates, *A. suum* L3 (~500) were exposed to 50,100 and 200 µM of the respective drugs: **(a)** thiabendazole, **(b)** artemisinin, **(c)** artesunate, **(d)** artemether for 24 and 48 hours and the metabolic activity assessed using the resazurin reduction assay. Resazurin was added at a final concentration of 7.5 µg/ml in each well either four or 24 hours after drug exposure for 24- and 48-hour exposure experiments, respectively. The relative fluorescence intensities were measured at 540 nm Ex/590 nm Em using a microplate reader (Synergy H1, BioTek Germany). Data represented is of two independent assays done in quadruplicate (24 hour) and three independent assays done in triplicate (48 hour) and normalized as % relative to DMSO. Whiskers indicate 95% percentile. Significance is represented by $p<0.05$: *, $p<0.005$: **, $p<0.0005$: ***, $p<0.00005$: ****. Dunnett's test used to compare each drug concentration to DMSO control at the same time.

**Supplementary Fig. S5**

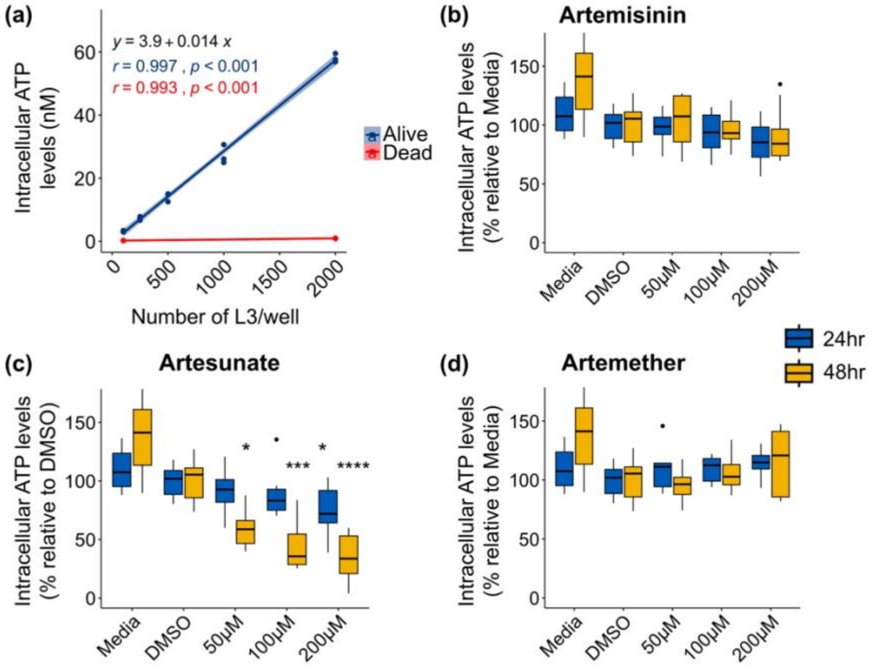


**Fig. S5. Exposure to artesunate decreases intracellular ATP levels of *A. suum* L3.** The ATP bioluminescence assay was optimized for its capacity to detect intracellular ATP content of live *A. suum* L3 as well as its sensitivity to sample size. In 96 well plates, increasing samples sizes of live and heat killed *A. suum* L3 were used to determine the capacity of the assay to detect ATP effectively **(a)**. *A. suum* L3 (~400) were exposed to 50, 100 and 200 μM of **(b)** artemisinin, **(c)** artesunate, **(d)** artemether for 24 and 48 hours and intracellular ATP levels assessed using the Promega CellTiter-Glo® 3D Cell Viability Assay kit following the manufacturer's instructions with slight modifications. DMSO was used as the vehicle control for the drugs. Data represented is of two independent assays done in quadruplicate and normalized as % relative to respective control. Whiskers indicate 95% percentiles. Significance is represented by $p<0.05$: *, $p<0.0005$: ***. Dunnett's' test used to compare each drug concentration to DMSO.

### Supplementary Fig. S6

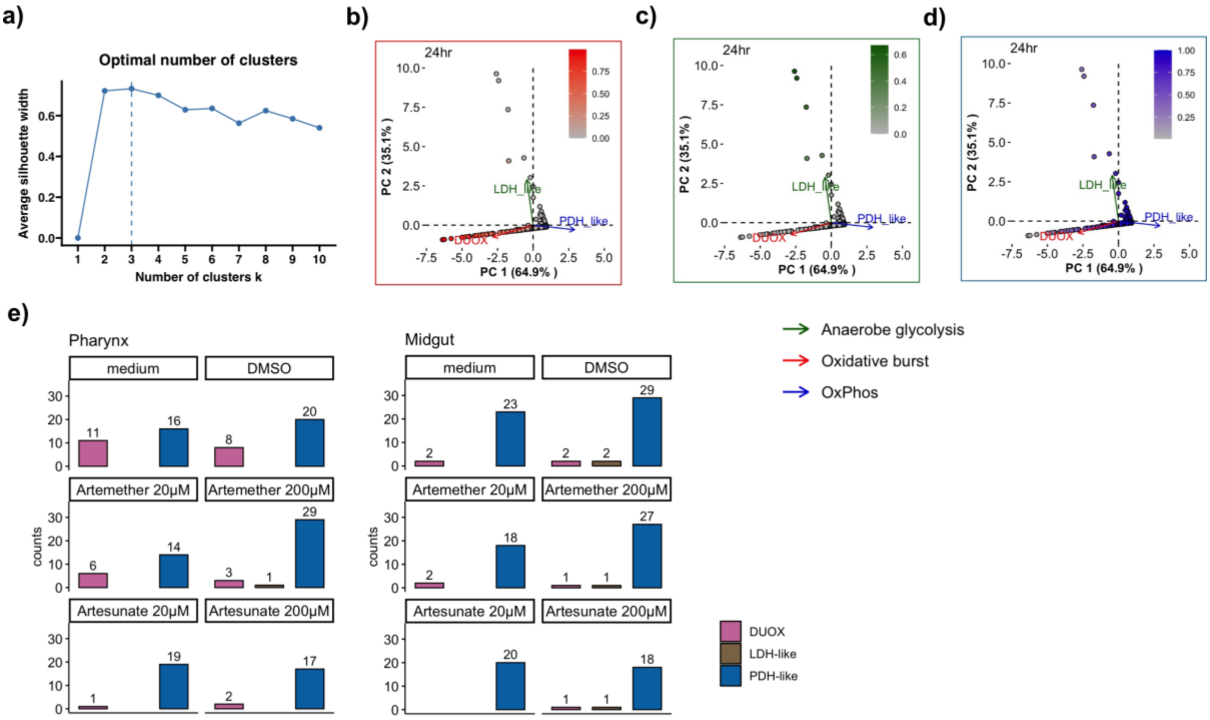


**Fig. S6. K-means clustering and metabolic profile identification.** Graph **(a)** illustrates the pre-determined optimal number of clusters determined by the silhouette method. The PCA biplots indicating the larvae associated with DUOX (**b**)**,** LDH-like **(c),** and PDH-like **(d)** metabolic profiles within PC1 and PC2.The gradients grey-green, grey-blue and grey-red represent increasing intensity from absence to presence of LDH-like, PDH-like, and DUOX-like metabolic profiles respectively. Bar plots **(e)** illustrating distribution of metabolic profiles within the pharynx and midgut.

**Supplementary Fig. S7**

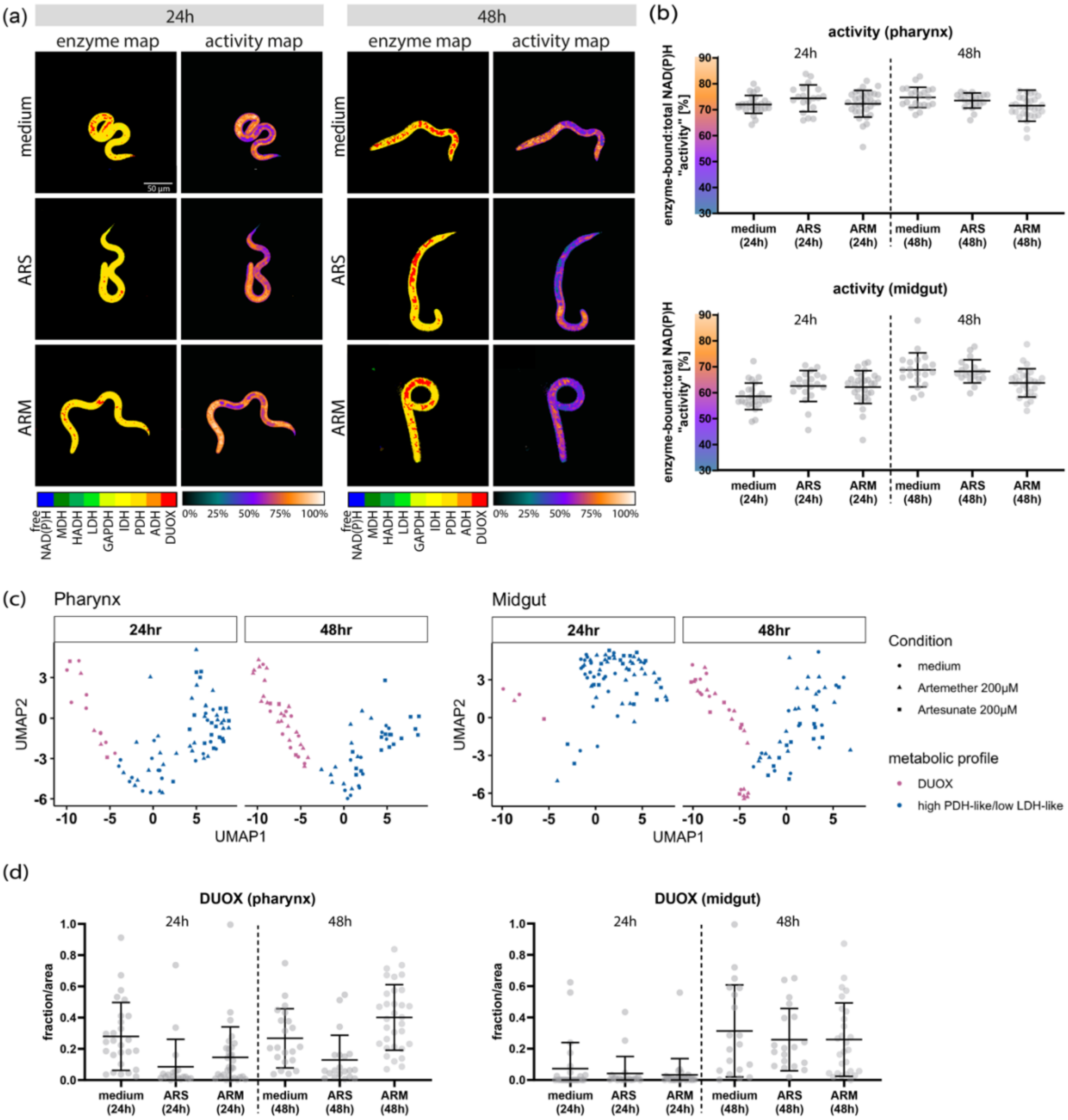


**Fig. S7.** ***A. suum*** **larvae exposed to ARTs for 48 hours in comparison to 24 hours.** *A. suum* L3 were exposed to 200 µM of artesunate (ARTS) and artemether (ARTM) for 48 hours and evaluated using two-photon NAD(P)H-FLIM. A representative microscopic image of L3 **(a)** visually illustrating the NAD(P)H enzyme map and activity map in 24-and 48-hour ART-exposed larvae. Dot plots **(b)** depicting metabolic activity within the pharynx and midgut at 24- and 48-hour timepoints. The UMAP plot **(c)** illustrating distribution of DUOX enzyme activity (depicted by magenta-pink) and high PDH-like/ low LDH-like enzyme activity (depicted by blue) in the pharynx and midgut at 24- and 48-hour timepoints. Dot plots **(d)** depicting DUOX enzyme activity within the pharynx and midgut.
